# Analysis of the study of the cerebellar pinceau by Korn and Axelrad

**DOI:** 10.1101/001123

**Authors:** Antonin Blot, Boris Barbour

## Abstract

The axon initial segment of each cerebellar Purkinje cell is ensheathed by basket cell axons in a structure called the pinceau, which is largely devoid of chemical synapses and gap junctions. These facts and ultrastructural similarities with the axon cap of the teleost Mauthner cell led to the conjecture that the pinceau mediates *ephaptic* (via the extracellular field) inhibition. Korn and Axelrad published a study in 1980 in which they reported confirmation of this conjecture. We have analysed their results and show that most are likely to be explained by an artefactual signal arising from the massive stimulation of parallel fibres they employed. We reproduce their experiments and confirm that all of their results are consistent with this artefact. Their data therefore provide no evidence regarding the operation of the pinceau.

## 1 Introduction

In the cerebellar cortex, multiple basket cell axon terminals enlace Purkinje cell somata, forming ‘baskets’ containing chemical synapses. The axons then extend to wrap around the initial segment of the Purkinje cell axon, creating a structure called the pinceau (Ramón y Cajal, 1911), which is largely devoid of chemical and electrical synapses (Sotelo and Llinás, 1972; Bobik et al., 2004; Iwakura et al., 2012). The basket cell axons are linked by septate junctions (Sotelo and Llinás, 1972). These properties led several groups to highlight a potential analogy with the Mauthner cell axon cap (Palay, 1964; Fox et al., 1967; Sotelo and Llinás, 1972), a much larger structure in fish that mediates ephaptic inhibition via the electrical field surrounding the axon initial segment (Furukawa and Furshpan, 1963; Furshpan and Furukawa, 1962).

Korn and Axelrad (1980) reported a study of the pinceau using very intense stimulation of parallel fibre bundles to excite basket cells indirectly. Early intracellular responses in Purkinje cells were attributed to the action of the pinceau, as was an early inhibition of antidromic spikes. We analysed their results and reproduced some of their experiments in cerebellar slices. We found that the signals recorded in the Purkinje cell with the same experimental arrangement are generated by the parallel fibre volley and not by the pinceau. Further analysis showed that all of the results that Korn and Axelrad attributed to the pinceau could be explained by this artefact.

## 2 Methods

### 2.1 Slice preparation

Animal experimentation methods complied with French and European regulations. Cerebellar slices were prepared from adult C57BL/6 female mice (*>* 8 weeks, Janvier or Charles River). Mice were anæsthetised with isoflurane (Nicholas Piramal India Ltd.) and killed by decapitation. The cerebellum was rapidly dissected into a cold solution containing the following (in mM) (Dugué et al., 2009): 130 K-gluconate, 15 KCl, 0.05 EGTA, 20 HEPES, and 25 glucose, with pH adjusted to 7.4 by NaOH, bubbled with 95% O_2_/5% CO_2_ and supplemented with 50 *µ*M D-APV. Sagittal slices (360 *µ*m) were cut in the same solution, using a Campden Instruments 7000smz slicer and stored at 32 °C in standard extracellular saline (bicarbonate-buffered solution; BBS), containing (in mM): 135 NaCl, 26 NaHCO_3_, 3 KCl, 1.25 NaH_2_PO_4_, 2 CaCl_2_, 1 MgCl_2_ and 25 D-glucose, bubbled with 95% O_2_/5% CO_2_.

### 2.2 Recordings

Recordings were performed at 32°C in BBS under a BX51WI microscope (Olympus) equipped with a CoolSNAP EZ camera (Photometrics) controlled using microManager (Edelstein et al., 2010) and ImageJ (Abramoff et al., 2004). Whole-cell current-clamp recordings were obtained using a Multiclamp 700B (Molecular Devices) with bridge balance and capacitance neutralisation.

Experiments were controlled using the WinWCP freeware (John Dempster, Strathclyde Electrophysiology Software). The composition of the pipette solution was (in mM): 0.4 Na-GTP, 0.5 L-(−)-Malic acid, 0.008 Oxaloacetic acid, 0.18 *α*-Ketoglutaric acid, 0.2 Pyridoxal 5’-phosphate hydrate, 5 L-Alanine, 0.15 Pyruvic acid, 15 L-Glutamine, 4 L-Asparagine, 1 L-Glutathione reduced, 10 Hepes, 4 KCl, 10 GABA, 2.1 Mg-ATP, 1.4 Na-ATP, 5 Phospho-creatine-K_2_, 0.5 K_3_-Citrate, 120 K-Gluconate, 0.1 EGTA, 2.2 K_2_-Phosphate, 0.05 CaCl_2_. We purchased: chemicals from Sigma; drugs from Ascent, Tocris and Sigma (TTX).

Potentials are reported without correction for junction potentials.

Extracellular stimulation with an isolated stimulator (model 2100, A-M Systems) employed patch pipettes (resistance 1–10 MΩ) filled with a HEPES buffer solution containing (in mM) 141 NaCl, 2.5 KCl, 1.25 NaH_2_PO_4_, 10 HEPES, 1.6 CaCl_2_ and 1.5 MgCl_2_ and 25 glucose. It was necessary to shield stimulator cables as well as the non-immersed portions of recording and stimulating electrodes to prevent stimulus artefacts from influencing the somatic voltage.

### 2.3 Data analysis

Electrophysiological data were analysed in Python using custom software relying on the *numpy* and *scipy* packages (Jones et al., 2001). Data are reported as mean ± sem. Non-parametric tests were preferred and performed in *GNU R* (R Development Core Team, 2011). Two-tailed tests were systematically used.

## 3 Results

### 3.1 Analysis of Korn and Axelrad (1980)

We first summarise the different experiments in the Korn and Axelrad (1980)^1^:

1. Basket cells were excited indirectly by very intense stimulation of parallel fibres (~ 1000 times greater currents than we used in the experiments shown below).
2. The authors present intracellular recordings in Purkinje cells of ‘passive hyperpolarising potentials’ (PHPs) that preceded chemical synaptic inhibition. These are illustrated in Figs. 1B1,4, 3A2–5 and 4B1,2. (In the latter two figures, the subtraction of the intracellular from the extracellular voltage was performed to estimate the true somatic membrane potential.)
3. Inhibition in Purkinje cells of antidromic spikes preceding chemical inhibition was shown by extracellular single unit (Fig. 1B2,3) and field (Fig. 2) recordings.
4. PHPs were shown to be insensitive to changes of the intracellular chloride concentration (Fig. 3).
5. The timing of extracellular spikes attributed to basket cells was analysed (Figs. 4A2, B2 and C).
6. A model of the inhibitory mechanism was proposed in Fig. 4D.

It is useful to begin by analysing the model. The basket cell action potential was conjectured (but not demonstrated) to produce a positivity in the confined extracellular space of the pinceau. In the model, this positive voltage (change) is transferred to the inside of the Purkinje cell axon via the resistance labelled ‘RIs’, the membrane resistance of the initial segment. In reality, the admittance of membrane at frequencies related to action potentials is likely to be dominated by the capacitance rather than the conductance^2^. Similar arguments apply to the ‘presynaptic’ membrane, but the capacitive current flowing out of the basket cell axon was also ignored.

Irrespective of the capacitive or resistive nature of the current entering the Purkinje cell axon, the inward current would be expected to *depolarise* the Purkinje cell soma. The PHP therefore has the opposite sign to that expected and does not represent a direct recording of the pinceau effect. This fact was appreciated by Korn and Axelrad, who state in their discussion: ‘…*the positivity is masked, however, by a predominantly negative field generated by active elements in the molecular layer*’. The action of the masking hyperpolarisation is presumably inhibitory, but would not reflect an action of the pinceau.

The next question concerns the nature of the masking event/PHP. The description of the model implies that it represents basket cell action potentials somewhere outside the pinceau (the authors propose the molecular layer, but one should also consider the axons forming the basket around the Purkinje cell soma). The Purkinje cell would essentially operate as an extracellular electrode, with an extracellular negativity resulting in an intracellular negativity. However, another possible driving potential is the extracellular field of the action potentials of the parallel fibre bundle (with the Purkinje cell again acting like an extracellular electrode). Korn and Axelrad argued that the parallel fibre volley had ‘*inappropriate timing*’ for this hypothesis because in Fig. 2 the onset of the extracellular volley, measured by its first positive peak, slightly preceded the early inhibition of the antidromic field. Nevertheless, the central negative peak, which would produce the intracellular hyperpolarisation, occurred 1 ms later (Fig. 4A1). Furthermore, the deep parallel fibre volley (shown in Fig. 4A2) was synchronous with the intracellularly recorded PHP in Fig. 1B3 and B4. The data therefore support the hypothesis that the field of the parallel fibre volley generated the PHP.

We note that the insensitivity of the PHP to the intracellular chloride concentration does not exclude the hypothesis that the PHP is generated by the parallel fibre volley, as no chloride conductance would be involved.

Two further arguments can be made in favour of the PHP being caused by the parallel fibre volley rather than basket cell action potentials in the molecular and/or Purkinje cell layers.

1. Fig. 4A1 shows that the extracellular field of the parallel fibre volley is much greater than that of basket cell action potentials (arrow Fig. 4A2) and the area of Purkinje cell dendrite exposed to the volley is probably much more extensive. This suggests that the parallel fibre volley will dominate.
2. The limited data shown on the timing and precision of basket cell spikes (Fig. 4C) suggest that the PHP begins somewhat *before* basket cell spikes. This would of course be incompatible with the proposed mechanism in which those spikes cause the PHP.

### 3.2 Experimental verification

We now turn to experiment to address the remaining questions. The above analysis strongly suggested that the PHP reflects the parallel fibre volley rather than basket cell action potentials. We therefore predicted that prevention of basket cell excitation should not abolish the PHP.

We recorded from Purkinje cells in adult mouse transverse cerebellar slices and stimulated a bundle of parallel fibres (Fig. 1). Preceding chemical synaptic excitation (occasionally absent; we stimulated just off beam in order to reproduce similar data to Korn and Axelrad) and inhibition, we recorded voltage transients with a form and timing compatible with the reported PHPs, but somewhat smaller (Fig. 1b,d, 211 ± 83**µ**V vs 1.14 ± 0.33 mV). This amplitude ratio is similar to that of the parallel fibre volley measured extracellularly, which was 418 ± 170 **µ**V in our hands (Fig. 1c) compared to more than 5 mV in Korn and Axelrad (Fig. 4A1). Our smaller intracellular and extracellular responses may reflect a reduced number of viable parallel fibres in slices, shunting of the extracellular field by the bath solution and/or our use of *much* smaller stimulation intensities (a few *µ*A compared to up to mA for Korn and Axelrad). Application of NBQX (5 *µ*M) should block excitation by parallel fibres of basket cells (and Purkinje cells). This was confirmed by the abolition of the EPSP and IPSP in the Purkinje cell. In accord with our hypothesis that the PHP reflects the parallel fibre volley, it was unaffected by NBQX (Fig. 1f; 178 ± 56 **µ**V, 93 ± 8% of control, *p* = 0.5 paired t-test, *n* = 4); it was however abolished by TTX (100 nM, Fig. 1h, 3 ± 2 **µ**V, 4 ± 2% of control, *p* = 0.002, paired t-test, *n* = 4). These experiments show that the PHP is caused by the parallel fibre action potential volley and is not related to basket cell activity.

**Figure 1.**
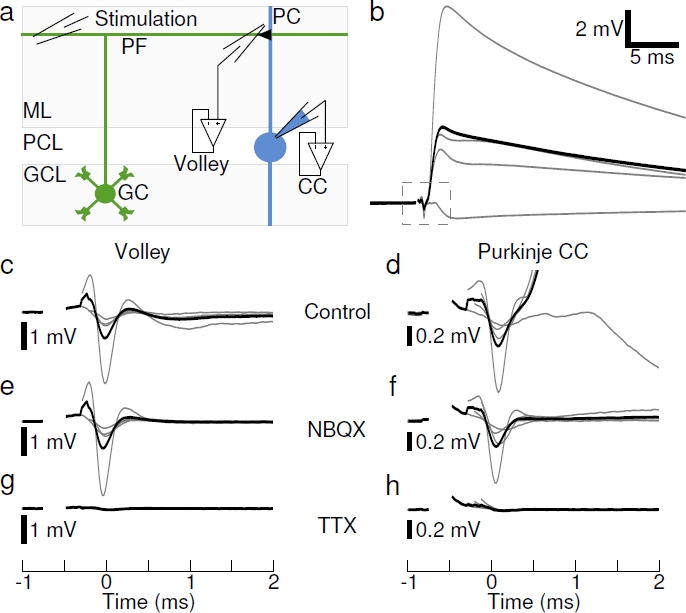
Passive hyperpolarisation. **a**, Recording configuration. Voltage responses to electrical stimulation in the molecular layer were recorded intracellularly in the Purkinje cell soma and extracellularly in the molecular layer. **b**, The stimulation induced a postsynaptic response in the Purkinje cell preceded by a fast and transient hyperpolarisation (centre of dashed box). The afferent volley recorded extracellularly **c** was synchronous with this intracellular hyperpolarisation, shown magnified in **d**. Both were unaffected by application of 5 *µ*M NBQX **e, f**, which abolished the postsynaptic potentials. The afferent volley and the intracellular hyperpolarisation were sensitive to 100 nM TTX **g**, **h**. All traces were aligned on the negative peak of the extracellular volley in control conditions. Stimulus artefacts were blanked.

Finally, although Korn and Axelrad probably did not directly record any signals arising from the pinceau, the question remains as to whether the inhibition of antidromic spikes in their Figs. 1 and 2 nevertheless reflected an effect of the pinceau or whether it could be explained by the parallel fibre volley. Both the single unit and field recordings in Korn and Axelrad are likely to represent action potential invasion of the axon initial segment and Purkinje cell soma(ta), as these compartments generate by far the largest part of the extracellular signal. We sought to test whether this invasion could be inhibited by a small hyperpolarisation of the size of the PHP in the absence of activation of the pinceau or whether some additional inhibitory mechanism was necessary.

**Figure 2.**
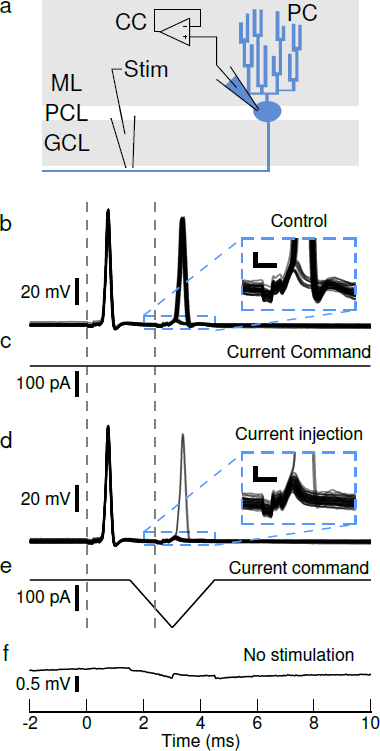
Inhibition of antidromic spikes. **a**, Recording configuration. Doublets of antidromic spikes elicited by white matter stimulation were recorded in current clamp in Purkinje cells maintained near -60 mV. On alternate sweeps, a small hyperpolarising current was injected in the Purkinje cell soma just before and during the second stimulus. The second of the antidromic spikes, induced at the end of the relative refractory period, invaded the soma in 72% of the trials **b** (and associated inset) in the absence **c** of the injected current and 8% of trials **d** when the current was injected **e**. **f**, Average hyperpolarisation induced by the current injection in the absence of electrical stimulation. **b**, **d**, Overlays of 36 sweeps; vertical dashed lines indicate electrical stimuli. Scale bar for insets: 2 mV, 0.5 ms. **c**, **e**, **f**, Averages of 28 sweeps.

We stimulated antidromic action potentials while recording intracellularly from Purkinje cells (Fig. 2). (Particular care was required to shield stimulator cables and electrodes to prevent capacitive coupling of the stimulating voltages to the recording electrode from exciting the Purkinje cell at the soma; Methods.) We found that invasion of the soma (and presumably the axon initial segment) by antidromic action potentials was very sensitive to voltage. We reproduced the experimental arrangement of Fig. 1 of Korn and Axelrad, in which a doublet of antidromic spikes were elicited and the interval between the two stimuli (less than 3 ms) was adjusted so that the second spike occurred at the end of the relative refractory period and exhibited some failures at the soma (Fig. 2b). A short injection of current inducing a hyperpolarisation of less than 1 mV (the size of the PHPs reported by Korn and Axelrad), applied just before and during the second antidromic stimulation (Fig. 2e, f), was sufficient to reduce significantly the number of somatic action potentials (62.2 ± 2.1% of the trials in control, 14.6 ± 1.9% with current injection, *p* = 0.008, paired Wilcoxon test, *n* = 8). Excitation at the site of stimulation in the axon was, however, not affected, as small all-or-none events could still be recorded when somatic invasion failed (Fig. 2d inset; no failures were observed during the recording shown). These small responses presumably reflected activity at one of the proximal nodes of Ranvier.

These experiments show that the small hyperpolarisation produced by the parallel fibre volley is able to prevent antidromic action potentials from invading the soma. The existence of such inhibition therefore does not require an additional inhibitory mechanism.

## 4 Conclusion

We conclude that the results of Korn and Axelrad could all be explained by the parallel fibre volley and therefore provide no evidence concerning the operation of the pinceau.

## 5 Acknowledgements

This work was supported by the ANR (ANR-08-SYSC-005, ANR-08-BLAN-0023) and the ENS (fellowship to A.B.).

## Footnotes

1 Note that references to figures in Korn and Axelrad’s paper will be in plain text with capital letter panel labels, while references to figures in the present paper will be blue-coloured links with lower case panel labels.

2 The admittance at 500 Hz of the membrane capacitance will be 2*πf C* = 2.4 mS cm^*−*2^, compared to a specific membrane conductance of 8.1 *µ*S cm^*−*2^ (using the value for Purkinje cells from Roth and Häusser, 2001).

